# Rho-kinase mediates the anorexigenic action of melanocortin by suppressing AMPK

**DOI:** 10.1101/677880

**Authors:** Sang Soo Kim, Won Min Hwang, Won-Mo Yang, Hyon Lee, Kyong Soo Park, Yossi Dagon, Young-Bum Kim

**Author notes:** These authors contributed equally to this work (Sang Soo Kim, Won Min Hwang, Won-Mo Yang). Correspondence: Young-Bum Kim, Ph.D. Division of Endocrinology, Diabetes and Metabolism, Beth Israel Deaconess Medical Center, 330 Brookline Avenue, Boston, MA 02215.

## Abstract

**Objective:** Melanocortin action is essential for the maintenance of energy homeostasis. However, knowledge of the signaling mechanism(s) that mediates the effect of melanocortin remains incomplete.

**Methods:** ROCK1 is a key regulator of energy balance in the hypothalamus. To explore the role of ROCK1 in the anorexigenic action of melanocortin, we deleted ROCK1 in MC4R neurons in mice. Next, we studied the metabolic effects of MC4R neuron-specific ROCK1-deficiency and following treatment with α-melanocyte-stimulating hormone (MSH).

**Results:** Here we show that α-MSH increases Rho-kinase 1 (ROCK1) activity in the hypothalamus. Deficiency of ROCK1 in MC4R-expressing neurons results in increased body weight in mice fed normal chow diet. This is likely due to increased food intake and decreased energy expenditure. Importantly, we find that ROCK1 activation in MC4R expressing neurons is required for melanocortin action, as evidenced by the fact that α– MSH’s ability to suppress food intake is impaired in MC4R neuron-specific ROCK1-deficient mice. To elucidate the mechanism by which ROCK1 mediates melanocortin action, we performed *in vitro* studies in hypothalamic cells expressing MC4R. We demonstrate that α–MSH promotes the physical interaction of ROCK1 and Gα_12_, and this results in suppression of AMPK activity.

**Conclusions:** Our study identifies ROCK1 as a novel mediator of melanocortin’s anorexigenic action and uncover a new MC4R→Gα_12_→ROCK1→AMPK signaling pathway. Targeting Rho-kinase in MC4R-expressing neurons could provide a new strategy to combat obesity and its related complications.

## INTRODUCTION

Obesity has reached pandemic proportions globally and is associated with serious health risks including type 2 diabetes mellitus, atherosclerotic disease, and cancer resulting in a higher rate of premature death (Collaboration, 2017, Global et al., 2016). Obesity arises from an imbalance in energy metabolism linked to caloric intake, physical inactivity, and genetic risk factors (Ahima, 2011). The brain plays a key role in the control of body fat accumulation and energy homeostasis (Ryan et al., 2012, Schwartz and Porte, 2005). Neurons in the hypothalamus expressing the melanocortin-4 receptor (MC4R), a G-protein-coupled receptor (GPCR), as well as the paraventricular hypothalamus (PVH), amygdala, DMH, and cortex are crucial for the regulation of feeding behavior, energy expenditure, glucose homeostasis and autonomic outflow (Krashes et al., 2016, Balthasar et al., 2005, Small et al., 2001, Shah et al., 2014, Rossi et al., 1998, Huszar et al., 1997). MC4R coordinates energy balance by integrating signals for satiety and hunger in response to α-MSH and Agouti-related protein, respectively, released from distinct neuronal populations in the arcuate nucleus (Garfield et al., 2015). Deletion of MC4R in mice leads to maturity onset hyperphagic obesity (Huszar et al., 1997), while in humans, genetic variants and mutations have been associated with common and severe forms of obesity (Calton et al., 2009, Hebebrand et al., 2010). Although the melanocortin network in the brain is critical for the maintenance of body weight homeostasis, and defects in this control system are implicated in the pathogenesis of obesity, the neural signaling components responsible for the MC4R-mediated regulation of food intake and energy expenditure are not fully understood.

Emerging data demonstrate that the Rho-kinase (ROCK) signaling pathway is implicated in the pathogenesis of metabolic-related disorders, including diabetes and cardiovascular disease (Kajikawa et al., 2014, Zhou et al., 2011, Huang et al., 2013a). Evidence from genetic models have revealed that ROCK1 is a novel regulator of glucose homeostasis and insulin resistance as well as *de novo* lipogenesis in peripheral tissue (Lee et al., 2009, Lee et al., 2014, Huang et al., 2013a). In particular, our work suggests that ROCK1 negatively regulates AMPK activation, as evidenced by the findings that AMPK activity increases in the absence of ROCK1 but decreases when ROCK1 is active (Huang et al., 2018). In addition, activation of hepatic ROCK1 is required for the control of cannabinoid-induced lipogenesis in which AMPK functions as a downstream mediator of ROCK1 (Huang et al., 2018). It has been reported that the inhibition of hypothalamic AMPK is necessary for leptin’s effects on food intake (Minokoshi et al., 2004), highlighting a key role for AMPK in energy balance (Claret et al., 2007). In line with these observations, experimental evidence that ROCK1 regulates food intake and body weight homeostasis by targeting hypothalamic leptin receptor signaling (Huang et al., 2012) further raises the question of whether the ROCK1-AMPK signaling axis plays an important role in the metabolic action of melanocortin on feeding behavior. In fact, nothing is known about the involvement of ROCK1 in central melanocortin signaling that is essential for metabolic energy homeostasis.

In this study, we investigated the physiological role of ROCK1 in MC4R-expressing neurons in regulating energy balance and further elucidated the neuronal signaling cascade underlying the metabolic effect of melanocortin on food intake. Here we show that ROCK1 activation in MC4R-expressing neurons is required for the anorexigenic action of melanocortin, which is mediated by the MC4R→Gα_12_→ROCK1→AMPK signaling pathway.

## RESULTS

### Deletion of ROCK1 in MC4R-expressing neurons in mice leads to obesity

To determine the physiological role of ROCK1 in MC4R-expressing neurons in energy metabolism, we generated mice lacking ROCK1 in MC4R-expressing neurons by mating ROCK1 floxed mice with MC4R-2A-Cre transgenic mice. Of note, the homozygous MC4R-Cre mice display an obese phenotype (not shown), so in all further experiments, we used only heterozygous mice. Body weights were measured weekly in wide-type (WT), MC4R-Cre, ROCK1^*loxP/loxP*^ and MC4R-Cre; ROCK1^*loxP/loxP*^ mice on a normal chow diet up to 24 weeks of age. As important negative controls, we noted that male MC4R-Cre and ROCK1^*loxP/loxP*^ mice had similar body weight compared to WT mice (Fig. 1A). However, male MC4R-Cre; ROCK1^*loxP/loxP*^ mice had significantly increased body weight compared with ROCK1^*loxP/loxP*^ mice (Fig. 1A). At 24 weeks of age, the average body weight of MC4R-Cre; ROCK1^*loxP/loxP*^ male mice was ~17% higher than ROCK1^*loxP/loxP*^ mice (45.1 ± 1.2g vs. 36.4 ± 2.3g, *P*<0.01). Because WT, MC4R-Cre and ROCK1^*loxP/loxP*^ mice showed no difference in body weight, ROCK1^*loxP/loxP*^ mice were used in further experiments as the control group. There was a similar pattern of increased body weight in female MC4R-Cre; ROCK1^*loxP/loxP*^ mice compared to ROCK1^*loxP/loxP*^ mice (~10% increase; 36.5±1.9 vs. 32.8±0.7, *P*<0.05) (Fig 1D). The increased body weight is likely a result of an increase in fat mass in both male and female mice (Fig. 1B and 1E). These data are further confirmed by increased fat depots weights (Fig. 1C and 1F). Together, our data suggest that deficiency of ROCK1 in MC4R neurons causes an impairment of energy metabolism that can lead to an increase in body fat.

**Figure 1.**
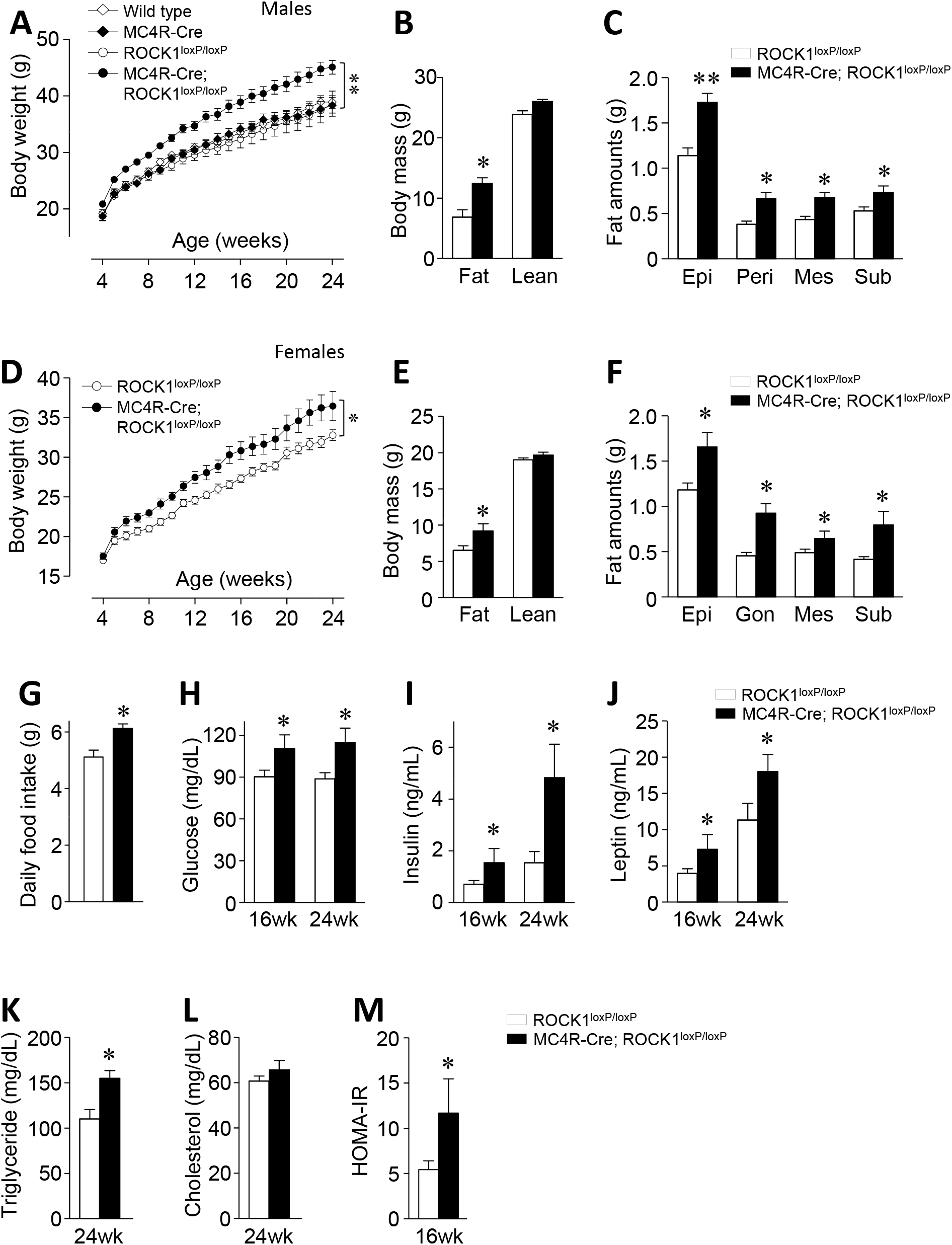
Mice lacking ROCK1 in MC4R-expressing neurons are obese. (A) Body weights (*n* = 8–12 per group), (B) body mass (*n* = 8–12 per group), and (C) fat depot (*n* = 11-13 per group) were measured in male wild type, MC4R-Cre, ROCK1^*loxP/loxP*^ and MC4R-Cre; ROCK1^*loxP/loxP*^ mice. (D) Body weights (*n* = 9–11 per group), (E) body mass (*n* = 9–11 per group), and (F) fat depot (*n* = 8–12 per group) were measured in female ROCK1^*loxP/loxP*^ and MC4R-Cre; ROCK1^*loxP/loxP*^ mice. (G) Daily food intake (*n* = 8 per group), (H) blood glucose (*n* = 7–8 per group), (I) serum insulin (*n* = 7–8 per group), (J) serum leptin (*n* = 4–8 per group), (K) serum triglyceride (*n* = 4–7 per group), (L) serum cholesterol (*n* = 5–7 per group), and (M) HOMA-IR (*n* = 7–8 per group) were measured in male ROCK1^*loxP/loxP*^ and MC4R-Cre; ROCK1^*loxP/loxP*^ mice. Body mass was measured by an MRI at 16 weeks of age. Blood glucose levels were measured from random fed mice. Serum parameters were measured from overnight fasted mice at 16 or 24 weeks of age. Data are presented as means ± SEM. * *P* < 0.05, ** *P* < 0.01 vs. ROCK1^*loxP/loxP*^ mice (control).

### Mice lacking ROCK1 in MC4R neurons display obesity-related metabolic changes

We further characterized the metabolic phenotype of MC4R neuron-specific ROCK1-deficient mice. Daily food intake in mice lacking ROCK1 in MC4R neurons significantly increased compared with control mice under chow-diet feeding (Fig. 1G). Thus, it appears that hyperphagia can explain the increased adiposity in these mice, suggesting that ROCK1 plays a pivotal role in regulating food intake. At 16 and 24 weeks of age, blood glucose, serum insulin, serum leptin, and serum TG levels significantly increased in MC4R neuron-specific ROCK1-deficient mice compared with control mice (Fig. 1H–K). However, serum cholesterol level was similar between the two groups (Fig. 1L). In addition, mice lacking ROCK1 in MC4R neurons are insulin resistant as evidenced by increased HOMA-IR (Fig. 1M) as well as ITT and GTT results (Supplementary Fig. 1A–B). These data demonstrate that genetic deletion of ROCK1 in MC4R neurons results in obesity-related metabolic disorders.

### ROCK1 deletion in MC4R-expressing neurons leads to decreased energy expenditure

To determine whether increased adiposity in MC4R neuron-specific ROCK1-dificient mice is due to decreased energy expenditure, oxygen consumption (VO_2_) and carbon oxide production (VCO_2_) were measured by the Comprehensive Lab Animal Monitoring System (CLAMS). Mice lacking ROCK1 in MC4R neurons had significantly lower VO_2_ consumption during light and dark cycles compared with control mice (Fig. 2A). Consistent with this, VCO_2_ was also greatly decreased in MC4R neuron-specific ROCK1-dificient mice (Fig. 2B). However, the respiratory exchange ratio (RER) was not different between the two groups (Supplementary data Fig. 2A), suggesting that the relative contribution of carbohydrates and fatty acids to overall energy expenditure are similar. Interestingly, the physical activity of mice lacking ROCK1 in MC4R-expressing neurons was reduced compared with control mice (Supplementary data Fig. 2B). Collectively, our data demonstrate that the increased adiposity in MC4R neuron-specific ROCK1-deficient mice is likely due to decreased energy expenditure and physical activity.

**Figure 2.**
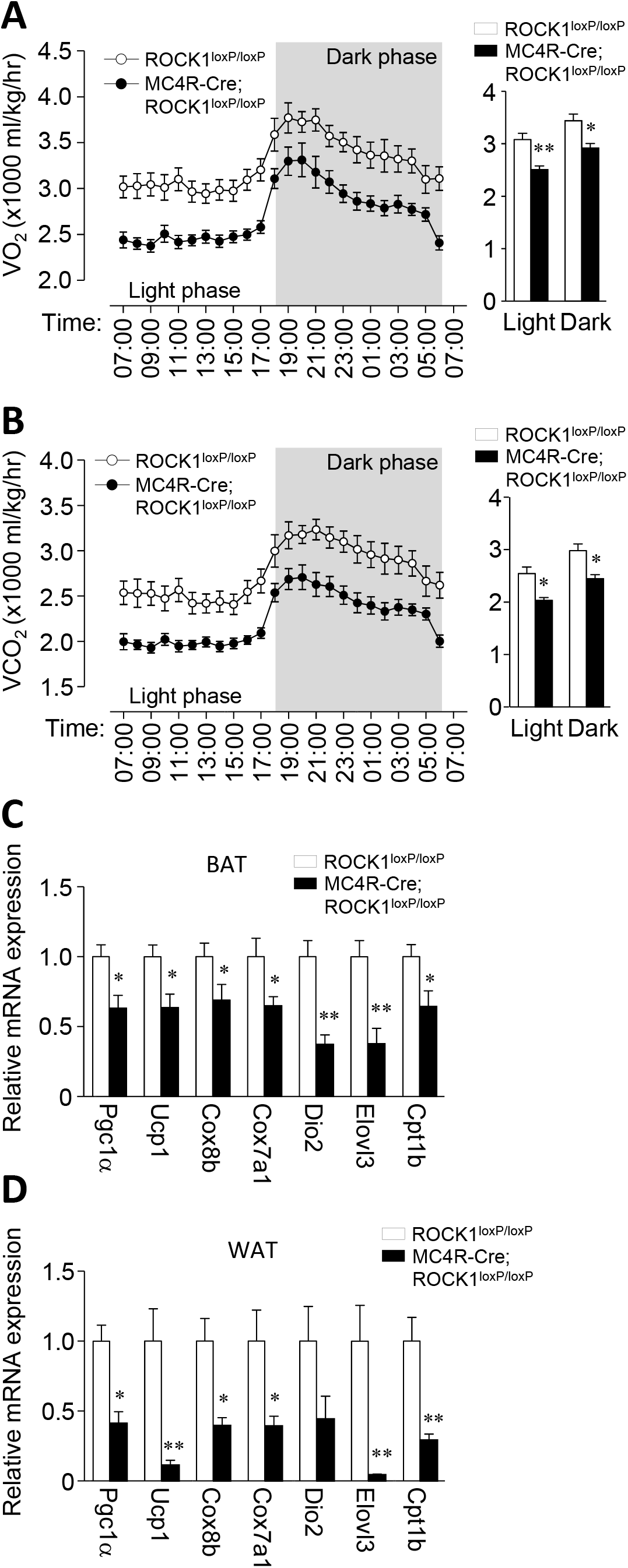
Deficiency of ROCK1 in MC4R-expressing neurons decreases energy expenditure. (A) O_2_ consumption (*n* = 8 per group), (B) CO_2_ production (*n* = 8 per group), (C) thermogenic gene expression in brown adipose tissue (BAT) (*n* = 8 per group) and (D) epididymal white adipose tissue (WAT) (*n* = 8 per group) were measured in male ROCK1^*loxP/loxP*^ and MC4R-Cre; ROCK1^*loxP/loxP*^ mice. Hourly average O_2_ consumption or CO_2_ production and corresponding light and dark phase oxygen consumption (12 h average) were assessed by CLAMS at 16 weeks of age. Thermogenic genes expression was measured from overnight fasted mice at 16 weeks of age (*n* = 5–7 per group). Data are presented as means ± SEM. * *P* < 0.05, ** *P* < 0.01 vs. ROCK1^*loxP/loxP*^ mice (control).

We further explored the mechanism by which ROCK1 deletion in MC4R neurons decreases energy expenditure by determining the thermogenic gene expression in brown adipose tissue (BAT) and white adipose tissue (WAT). Importantly, deletion of ROCK1 in MC4R neurons significantly decreased mRNA levels of thermogenic genes in BAT and WAT, including PGC1α, UCP1, Cox8b, Cox7a1, Elovl3, and Cpt1b (Fig. 2C–D). These data demonstrate that the decreased energy expenditure caused by ROCK1 deficiency in MC4R neurons could be explained, least in part, by the down-regulation of thermogenic gene expression.

### ROCK1 deletion impairs melanocortin action in MC4R neurons

To determine whether deficiency of ROCK1 in MC4R neurons alters the metabolic action of melanocortin on feeding behavior, we measured food intake in response to melanocortin receptor agonists α-MSH and MTII in ROCK1^*loxP/loxP*^ and MC4R-Cre; ROCK1^*loxP/loxP*^ mice. Fasting-induced food intake was increased in MC4R-Cre; ROCK1^*loxP/loxP*^ mice compared to ROCK1^*loxP/loxP*^ mice after 24 hours (Fig. 3A). Food intake in ROCK1^*loxP/loxP*^ mice markedly decreased 2–24 hours after the injection of intracerebroventricular (ICV) α-MSH and 2–8 hours after the injection of intraperitoneal (IP) MTII. However, the ability of α-MSH or MTII to suppress food intake in MC4R-Cre; ROCK1^*loxP/loxP*^ mice was impaired (Fig. 3B-C). ROCK1 deletion in MC4R-expressing neurons had no effect on neuropeptide gene expression, including POMC, AgRP, and NPY (Supplementary Fig. 3). Together, our data suggest that ROCK1 activation in MC4R neurons is required for the anorexigenic effect of melanocortin, and further indicate that the increased adiposity and caloric intake of MC4R neuron-specific ROCK1-deficient mice are caused by defective melanocortin signaling in MC4R neurons.

**Figure 3.**
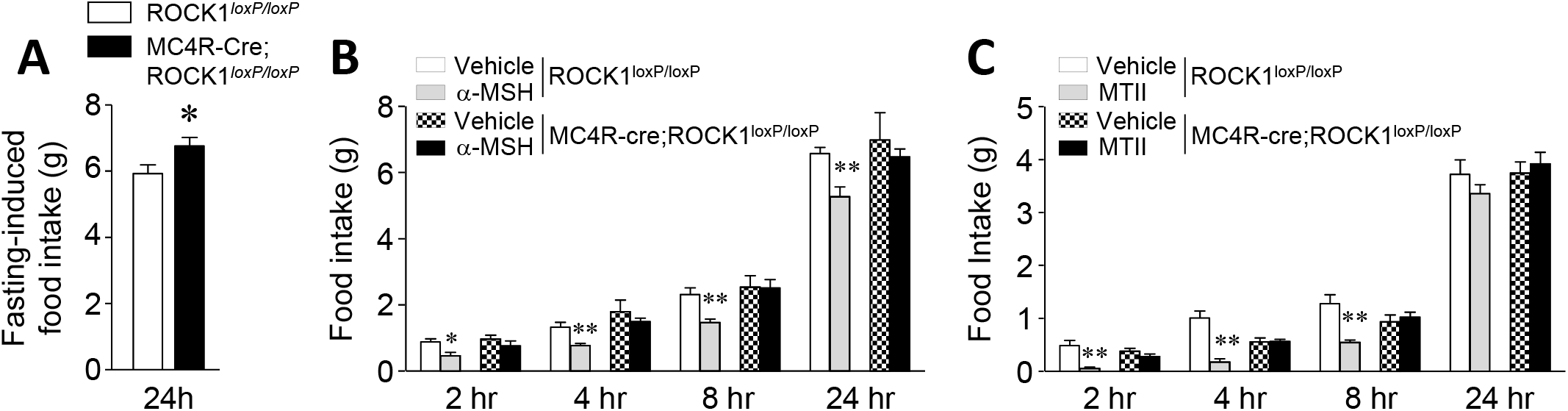
Anorexigenic effect of α-MSH is impaired in mice lacking ROCK1 in MC4R-expressing neurons. (A) Fasting-induced food intake (*n* = 8 per group), (B) α-MSH-induced food intake (*n* = 7–10 per group), and (C) MTII-induced food intake (*n* = 9–12 per group) were measured male ROCK1^*loxP/loxP*^ and MC4R-Cre; ROCK1^*loxP/loxP*^ mice. Fasting-induced food intake was measured from overnight fasted mice at 12 weeks of age. α-MSH-induced and MTII-induced food intake was measured 2, 4, 8 and 24 hours after ICV injections of α–MSH or IP injections of MTII at 16 weeks of age. Data are presented as means ± SEM. * *P* < 0.05, vs. ROCK1^*loxP/loxP*^ mice (control), * *P* < 0.01 vs. vehicle of ROCK1^*loxP/loxP*^ mice (control).

### Mice lacking ROCK2 in MC4R-expressing neurons are normal

To determine the physiological role of ROCK2 in energy balance, we generated mice lacking ROCK2 in MC4R-expressing neurons by mating ROCK2 floxed mice with MC4R-2A-Cre transgenic mice (Supplementary Fig. 4). Unlike MC4R-neuron-specific ROCK1-deficient mice, body weight and adiposity were indistinguishable between ROCK2^*loxP/loxP*^ and MC4R Cre; ROCK2^*loxP/loxP*^ mice on a normal chow diet (Fig. 4A–B). Food intake in MC4R-Cre; ROCK2^*loxP/loxP*^ mice was greatly decreased at 8 hours after IP injection of MTII, which was also observed in control mice (Fig. 4C). These data indicate that, in contrast to ROCK1, ROCK2 is unlikely to play a role in regulating energy homeostasis and feeding behavior.

**Figure 4.**
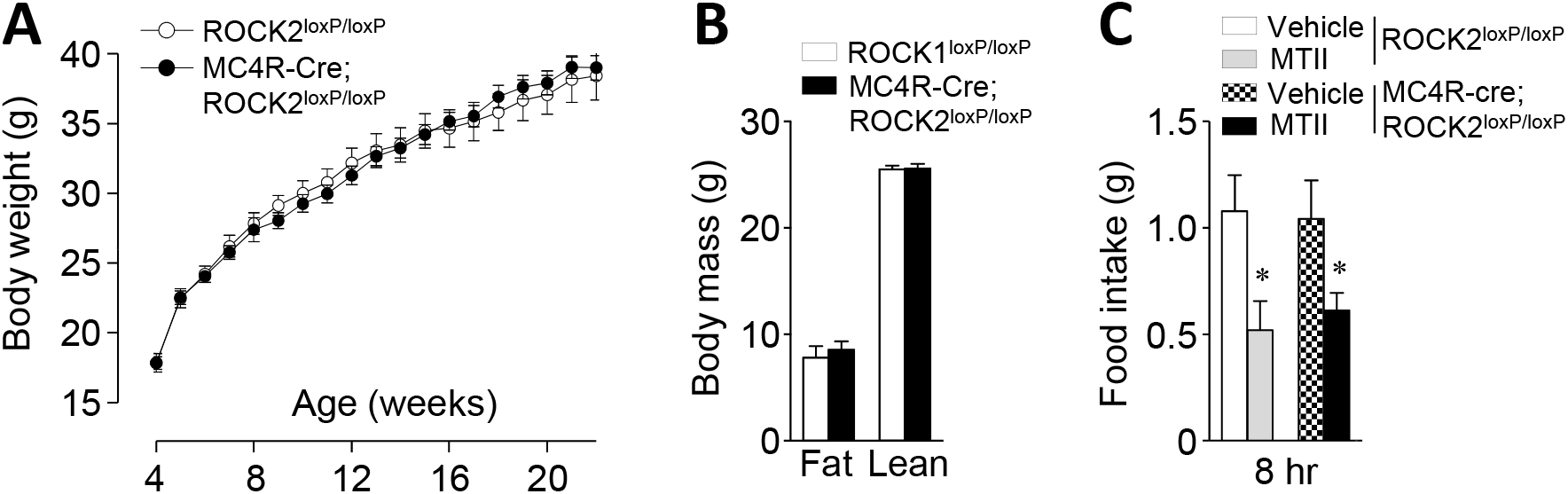
Mice lacking ROCK2 in MC4R-expressing neurons are normal. (A) Body weights (*n* = 12–16 per group), (B) fat mass (*n* = 12–16 per group), and (C) α-MSH-induced food intake (*n* = 5–8 per group) were measured in male ROCK2^*loxP/loxP*^ mice and MC4R-Cre; ROCK2^*loxP/loxP*^ mice. Body mass was measured by an MRI at 16 weeks of age. MTII-induced food intake was measured 2, 4, 8 and 24 hours after IP injections of MTII at 24 weeks of age. Data are presented as means ± SEM.

### α-MSH-induced cAMP production is mediated by MC4R

To investigate the MC4R signaling pathway, GT1-7 cells overexpressing MC4R-3×HA (GT1-7_MC4R_) were generated and studied. We confirmed that MC4R was overexpressed in GT1-7_MC4R_ cells, evidenced by increased MC4R expression in membrane fractions and increased HA expression on the cell surface, compared to GT1-7 cells (Fig. 5A–B). It is known that α-MSH increases cAMP production (Molden et al., 2015). In normal GT1-7 cells, α-MSH-induced cAMP production was not increased (Fig. 5C). However, as expected, α-MSH increased cAMP production in GT1-7_MC4R_ cells in a dose-dependent manner (Fig. 5D), suggesting that GT1-7_MC4R_ cells are functional. In addition, α-MSH-induced cAMP production was suppressed when MC4R signaling was blocked dose-dependently, indicating that α-MSH-stimulated cAMP production is mediated by MC4R (Fig. 5E). Moreover, α-MSH’s ability to increase cAMP levels was suppressed by treatment with a ROCK1 inhibitor in GT1-7_MC4R_ cells, highlighting a key role for ROCK1 in regulating the melanocortinergic network (Fig. 5F).

**Figure 5.**
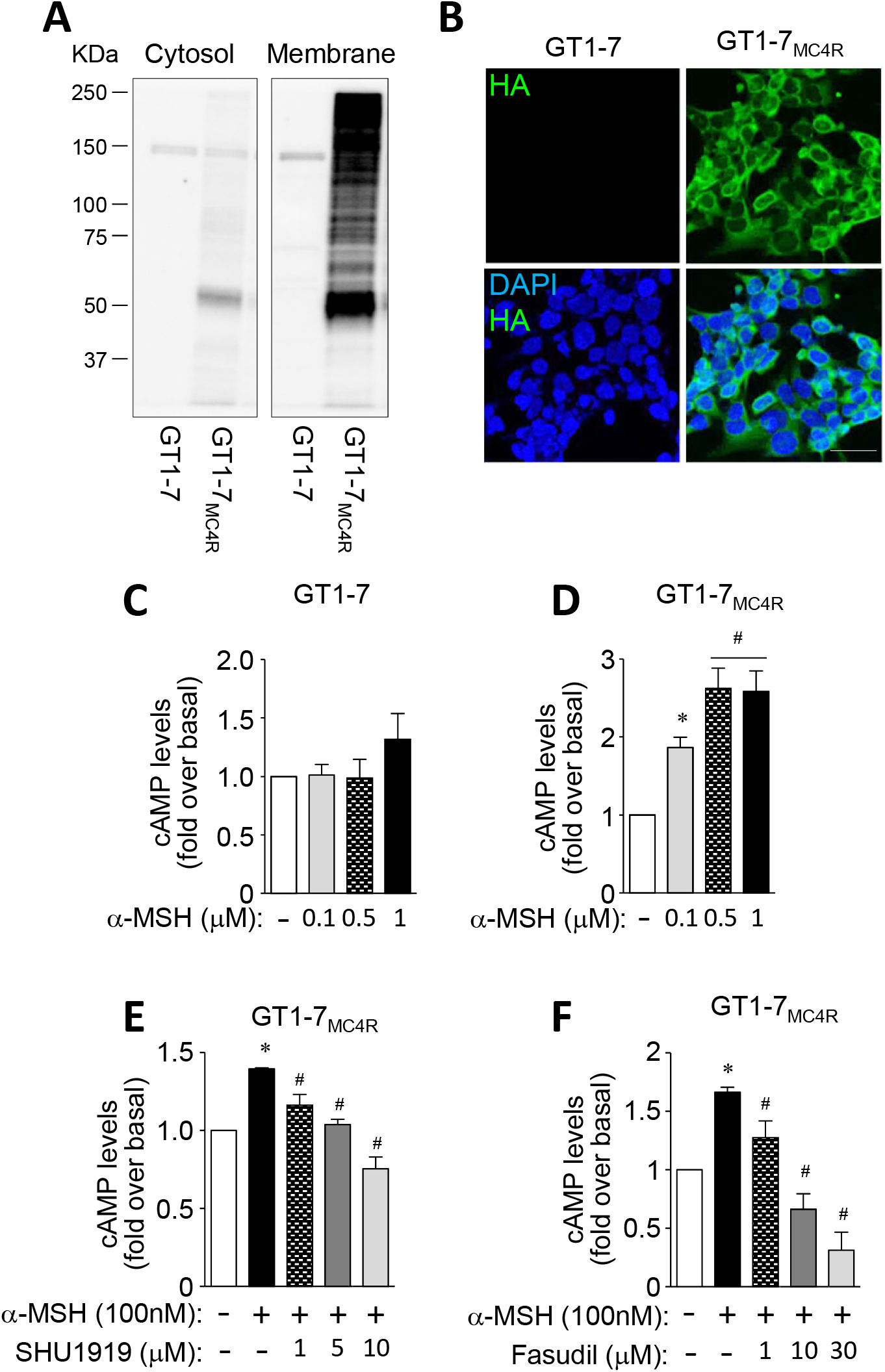
Generation of GT1-7 stable cell line overexpressing MC4R-3XHA (GT1-7_MC4R_). (A and B) Expression of MC4R was measured in GT1-7 cells and GT1-7_MC4R_ cells. (C and D) α–MSH-induced cAMP production was measured in GT1-7 cells and GT1-7_MC4R_ cells. (E) α– MSH-induced cAMP production was measured in the presence of SHU9119 in GT1-7 cells and GT1-7_MC4R_ cells. (F) α–MSH-induced cAMP production was measured in the presence of fasudil in GT1-7 cells and GT1-7_MC4R_ cells. MC4R protein levels from cytosol and membrane fractions were measured by immunoblotting analysis. Expression of MC4R in the cell membrane was measured by immunofluorescent analysis. Nuclei were visualized by DAPI staining (blue) and cell membranes were visualized by HA staining (green). Intracellular cAMP levels were measured by cAMP-Glo assay kit using a luminometer. Scale bar, 25 mm. Data are presented as means ± SEM and are representative of three independent experiments. * *P* < 0.05 vs. vehicle. ^#^*P* < 0.05 vs. α-MSH-treated cells.

### ROCK1 is a downstream mediator of the complexes of MC4R/Gα_12_ and MC4R/G_αs_

To establish the ROCK1 signaling pathway that mediates MC4R action on feeding, we measured the physical interaction of MC4R-to-Gα_12_ and Gα_12_-to-ROCK1 in GT1-7_MC4R_ cells using the *in situ* proximity ligation assay (PLA), a powerful tool for detecting protein interactions (Gullberg et al., 2004). α-MSH induced physical interactions between MC4R and Gα_12_ in GT1-7_MC4R_ cells (each red spot represents the detection of protein-protein interaction complexes), and this effect was abolished by treatment with the MC4R antagonist SHU9119 (Fig. 6A). However, inhibition of ROCK1 with fasudil had no effect on the interaction of the MC4R-to-Gα_12_ complex, suggesting that ROCK1 acts as a downstream signaling component of the complex of MC4R/Gα_12_. We also found that ROCK1 physically binds with Gα_12_ in response to α-MSH and that both ROCK1 inhibition and MC4R inhibition block this effect (Fig. 6B). Similar results were observed when we measured the physical interaction of MC4R-to-G_αS_ and G_αS_-to-ROCK1, i.e., α-MSH increased the interaction between MC4R and GαS, which occurred independent of ROCK1 inhibition, and between G_αS_ and ROCK1. α-MSH-induced interaction of G_αS_ and ROCK1 was suppressed by treatment with a MC4R and ROCK1 inhibitor (Fig. 6C–D). Collectively, these data demonstrate that ROCK1 acts as a downstream signaling component of the complexes MC4R/Gα_12_ and MC4R/G_αS_.

**Figure 6.**
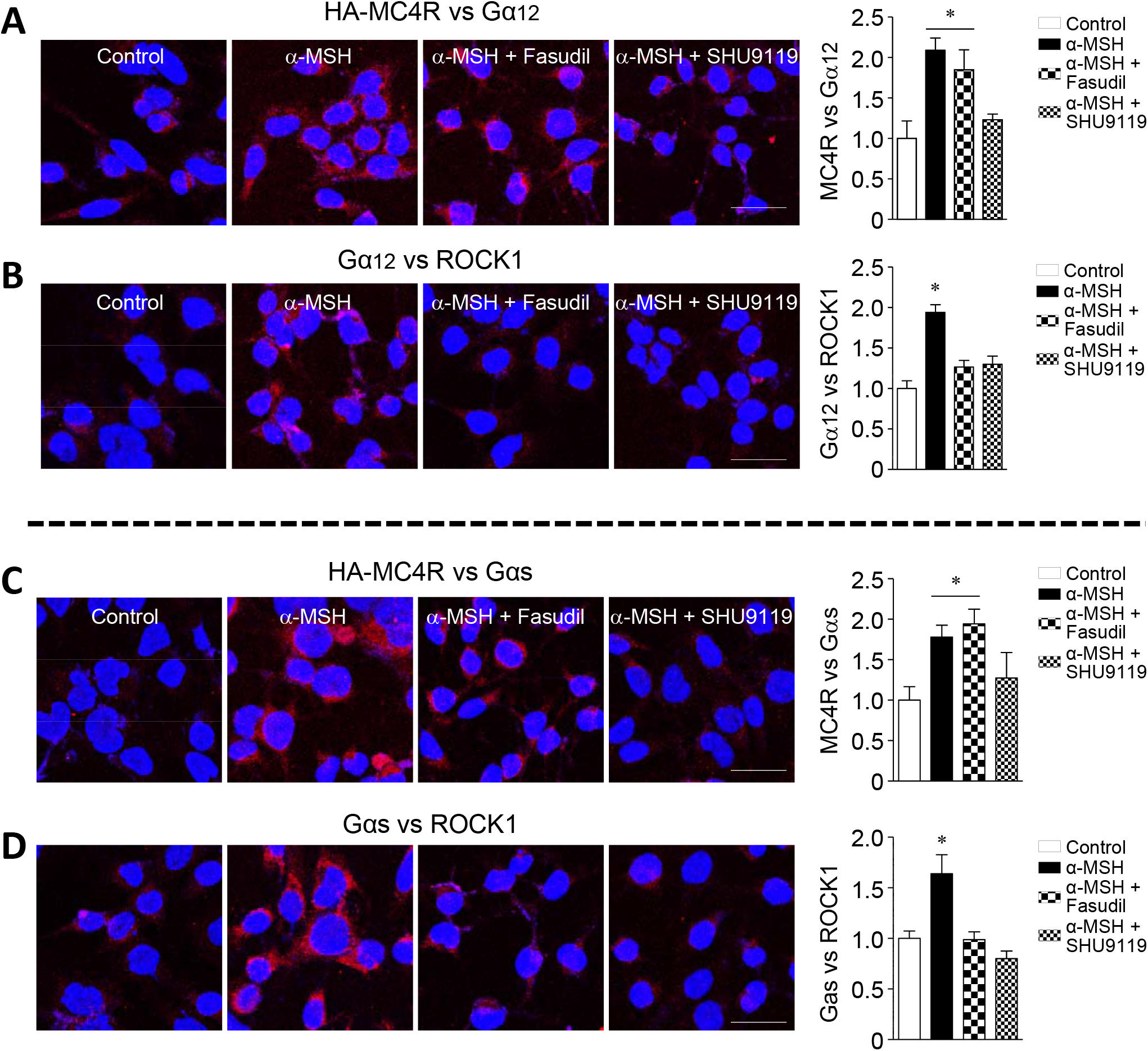
Melanocortin induces the physical interactions of MC4R-Gα_12_, ROCK1-Gα_12_, MC4R-G_αs_ and ROCK1-G_αs_. (A and B) Proximity ligation assays (PLA) of the interaction (red spots, blobs) between MC4R and Gα_12_ upon α–MSH treatment in the presence of fasudil or SHU9119 were performed in GT1-7_MC4R_ cells. (C and D) PLA of the interaction (red spots, blobs) between MC4R and Gαs upon α–MSH treatment in the presence of fasudil or SHU9119 were performed in GT1-7_MC4R_ cells. GT1-7_MC4R_ cells were treated as indicated. Each red spot represents MCR4-Gα_12_, ROCK1-Gα_12_, MCR4-Gαs or ROCK1-Gαs interactions. Bars show quantitation of all interactions as indicated. Nuclei were visualized by DAPI staining (blue). Scale bar, 12.5 mm. Data are presented as means ± SEM and are representative of three independent experiments. * *P* < 0.05 vs. control.

### ROCK1 mediates melanocortin action by suppressing AMPK

Hypothalamic AMPK has been implicated in the regulation of orexigenic and anorexigenic action by hormonal and nutritional signals (Minokoshi et al., 2004, Kahn et al., 2005). Thus, we tested the possibility that AMPK is involved in melanocortin’s anorexigenic action in the context of ROCK1 signaling. Treatment of GT1-7_MC4R_ cells with α-MSH led to a significant increase in ROCK1 activity but this effect was abolished by ROCK1 inhibition (Fig. 7A). α-MSH suppressed AMPK activity in GT1-7_MC4R_ cells; however, interestingly, AMPK activity was completely restored to control levels when ROCK1 was blocked (Fig. 7B), suggesting that ROCK1 negatively regulates AMPK activation. Consistent with our *in vitro* data, *in vivo* stimulation with α-MSH or MTII greatly increased hypothalamic ROCK1 activity in ROCK1^*loxP/loxP*^ mice (Fig 7C–D). However, as expected, ROCK1 activation did not respond to α-MSH or MTII in MC4R-Cre; ROCK1^*loxP/loxP*^ mice (Fig 7C and 7F). Concurrently, α-MSH or MTII-induced hypothalamic AMPK activity was significantly suppressed in ROCK1^*loxP/loxP*^ mice (Fig. 7D and 7G) but unaltered in MC4R-Cre; ROCK1^*loxP/loxP*^ mice (Fig. 7E and 7H). Together, these data clearly suggest that the activation of hypothalamic ROCK1 is required for the regulation of melanocortin-induced AMPK suppression that is crucial in regulating feeding behavior.

**Figure 6.**
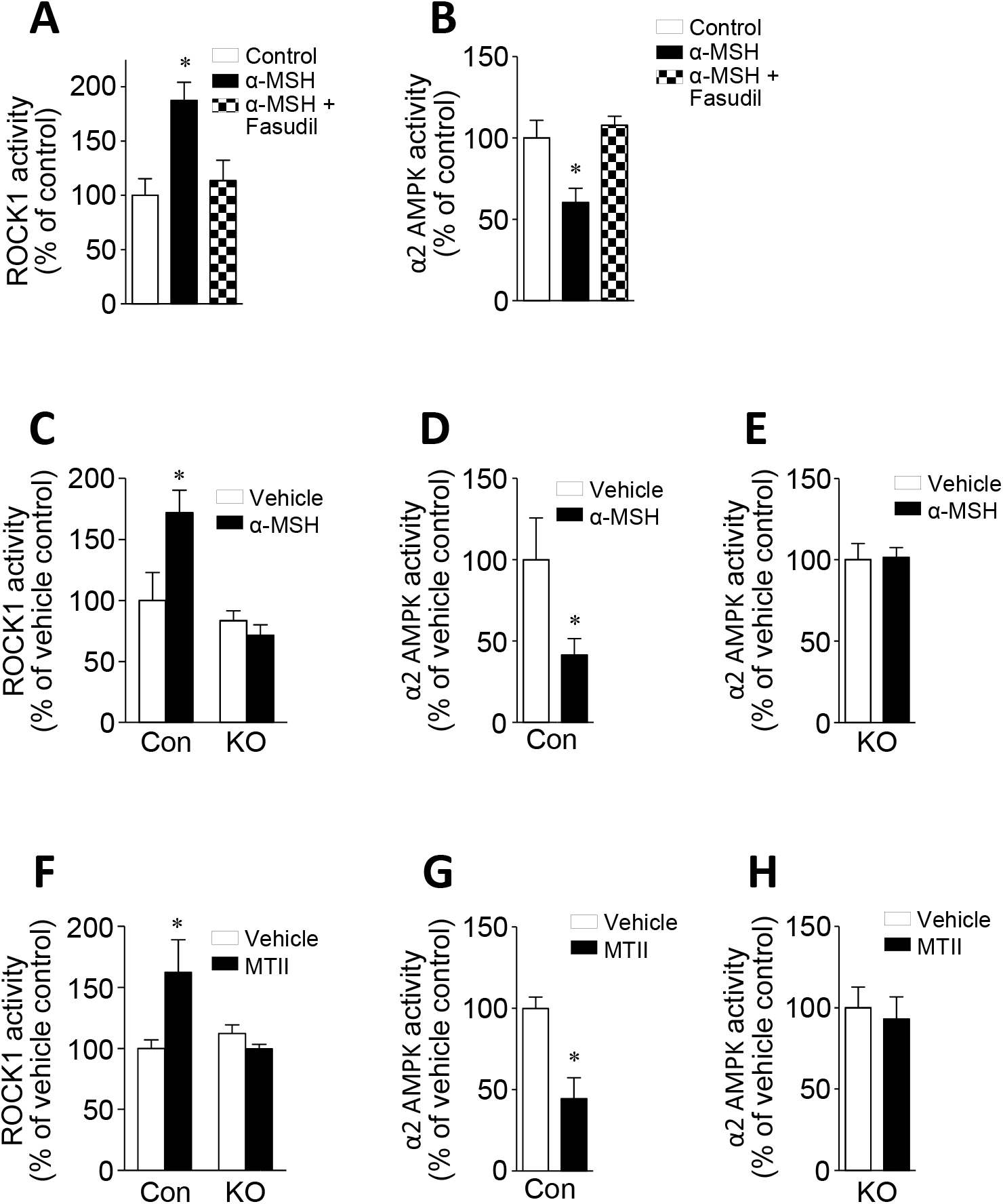
ROCK1 mediates melanocortin action by suppressing AMPK. (A) α-MSH-stimulated ROCK1 activity and (B) α-MSH-stimulated AMPK activity were measured in GT1-7_MC4R_ cells. GT1-7_MC4R_ cells were stimulated with α-MSH in the presence of fasudil. ROCK1 and AMPK activities were measured by immune-complex assay. Data are presented as means ± SEM and are representative of three independent experiments. * *P* < 0.05 vs. control. (C) α-MSH-stimulated ROCK1 activity was measured in the hypothalamus of male ROCK2^*loxP/loxP*^ mice and MC4R-Cre; ROCK2^*loxP/loxP*^ mice at 16 weeks of age (*n* = 3–6 per group). (D) α-MSH-stimulated AMPK activity was measured in the hypothalamus of male ROCK1^*loxP/loxP*^ mice (control) at 16 weeks of age (*n* = 5–8 per group). (E) α-MSH-stimulated AMPK activity was measured in the hypothalamus of male MC4R-Cre; ROCK1^*loxP/loxP*^ mice (*n* = 6–10 per group). (F) MTII-stimulated ROCK1 activity was measured in the hypothalamus of male ROCK2^*loxP/loxP*^ mice and MC4R-Cre; ROCK2^*loxP/loxP*^ mice at 16 weeks of age (*n* = 3–4 per group). (G) MTII-stimulated AMPK activity was measured in the hypothalamus of male ROCK1^*loxP/loxP*^ mice (control) at 16 weeks of age (*n* = 5–6 per group). (H) MTII-stimulated AMPK activity was measured in the hypothalamus of male MC4R-Cre; ROCK1^*loxP/loxP*^ mice at 16 weeks of age (*n* = 3–5 per group). ROCK1 and AMPK activity were measured after ICV injection of α-MSH or IP injection of MTII. Data are presented as means ± SEM. * *P* < 0.05 vs. vehicle.

## DISCUSSION

The melanocortin receptor is a key regulator of energy homeostasis, and its deficiency has been implicated in early-onset obesity and monogenic obesity (Oswal and Yeo, 2007, Farooqi et al., 2003). However, major gaps in our knowledge regarding the downstream signaling components involved in the central melanocortin network have been a limitation in the field. Here we show that activation of ROCK1 in MC4R-expressing neurons is necessary for the metabolic effects of melanocortin on food intake, which is regulated through the MC4R→Gα_12_→ROCK1→AMPK signaling axis. Thus, we identified ROCK1 as an important player of melanocortin’s anorexigenic action.

In this study, we investigated the role of ROCK1 in MC4R-expressing neurons in the regulation of energy balance. We found that under a normal chow diet, the deficiency of ROCK1 in MC4R-expressing neurons results in an increase in body weight, which is most likely due to increased food intake and decreased energy expenditure. The metabolic phenotypes of MC4R neuron-specific ROCK1-deficient mice are similar to those of *MC4R*^-/-^ and *POMC* deficient mice in the context of energy balance (Yaswen et al., 1999, Marsh et al., 1999), further highlighting a key role for ROCK1 in body-weight homeostasis. Given the fact that MC4R is expressed in a number of neurons in the brain, including the paraventricular hypothalamus (PVH), amygdala, DMH, and cortex (Krashes et al., 2016), it is important to know which neuronal sites are responsible for the metabolic phenotype of the MC4R neuron-specific ROCK1-deficient mice. In this regard, it has reported that mice homozygous for the loxTB MC4R allele, which prevents normal gene transcription and thus do not express MC4Rs, and are markedly obese (Balthasar et al., 2005). Importantly, using Sim1-Cre transgenic mice to restore MC4R expression in the PVH and a subpopulation of neurons in the amygdala, reduced adiposity by 60%. This effect occurred by normalizing increased food intake. However, the reduction in energy expenditure was unaffected, suggesting a functional divergence between energy intake and energy expenditure in the melanocortinergic neuronal pathways. In this regard, our preliminary data demonstrate that Sim-Cre; ROCK1^*loxP/loxP*^ mice also display an obese phenotype (not shown). It is therefore likely that the metabolic effect of ROCK1 may act through MC4R-expressing neurons in either the PVH, amygdala or both.

Hypothalamic AMPK activation plays a critical role in nutrient-derived appetite signaling and energy balance (Minokoshi et al., 2004). Studies have revealed that melanocortin inhibits AMPK activity in both the murine hypothalamus and hypothalamic cell lines (Minokoshi et al., 2004, Damm et al., 2012). Yet until now, the upstream signaling mediators for the melanocortin-induced AMPK inhibition have been elusive. Here we demonstrate that α-MSH suppresses AMPK activity in GT1-7_MC4R_ cells, while it increases ROCK1 activity and inhibition of ROCK1 restored the α-MSH-induced suppression of AMPK to control levels. Importantly, ROCK1 deficiency also impaired α-MSH’s ability to suppress food intake, suggesting that ROCK1 negatively regulates AMPK as an upstream effector in MC4R-expressing neurons. Support for this comes from our recent work showing that the activation of hepatic ROCK1 suppresses AMPK activity leading to lipid accumulation in the liver, whereas the deletion of hepatic ROCK1 stimulates AMPK activity preventing the development of hepatic steatosis (Huang et al., 2018). Collectively, these data suggest that ROCK1 is required for the regulation of normal melanocortin-induced anorexigenic action, and engages with AMPK. However, the underlying mechanism of how ROCK1 regulates AMPK activity remains to be elucidated. ROCK1 may not directly regulate AMPK as ROCK1 did not interact with AMPK (not shown).

It is evident that MC4R signaling involves a cascade of events initiated by α-MSH binding to MC4R, followed by activation of the Gαs protein, which results in increased cAMP levels (Yang et al., 1999, Gantz et al., 1993). cAMP has been shown to modulate cellular excitability and neuronal function in the regulation of adiposity (McKnight et al., 1998). Notably, our data suggest that ROCK1 activation is required for melanocortin-induced intracellular cAMP production in MC4R-expressing neuronal cells, as evidenced by the fact that α-MSH-stimulated cAMP production is decreased when ROCK1 is inactive. This may be due to the novel physical interaction observed between ROCK1 and the Gαs subunit on PLA. In contrast to the current study, a recent work demonstrated that the regulation of the firing activity of hypothalamic PVN neurons by α-MSH can be mediated independently of Gαs signaling by the ligand-induced coupling of MC4R to the closure of inwardly rectifying potassium channels, Kir7.1 (Ghamari-Langroudi et al., 2015). Others have shown that MC4R binding induces Gq-dependent phospholipase-C activity (Newman et al., 2006) as well as G-protein-dependent MAPK signaling (Daniels et al., 2003). Thus, MC4R may couple with multiple intracellular mediators to exert its physiologic effects.

Various signaling pathways appear to mediate the metabolic effects of melanocortin. Because Rho/Rho-kinase signaling has been shown to be activated via the Gα_12_ subunit of the G protein coupled receptor pathway (Vogt et al., 2003, Schmandke et al., 2007), in this study, we propose a hypothesis for the involvement of ROCK1 in MC4R-mediated Gα_12_ signaling triggered by α-MSH stimulation. Our data demonstrated that MC4R physically interacted with the Gα_12_ subunit, and then Gα_12_ bound to ROCK1 in response to α-MSH. The interaction of ROCK1 to Gα_12_ is a downstream signaling event of the complex of MC4R/Gα_12_ following α-MSH stimulation. Given that α-MSH increases ROCK1 activity but decreases AMPK activity, it is conceivable that increased Gα_12_-associated ROCK1 activity negatively regulates a downstream mediator of AMPK, which is an essential step in melanocortin action in MC4R-expressing neurons. As a result, the ability of α-MSH to stimulate anorexigenic action is promoted, leading to decreased feeding behavior. Thus, we have established a previously unknown signaling axis for melanocortin’s anorexigenic action.

Unlike the role of ROCK1 in the melanocortin system, ROCK2 deficiency in MC4R-expressing neurons is not responsible for the MC4R-regulated energy utilization observed in the current study. ROCK2 (also known as ROKα) shares 65% overall amino acid homology with ROCK1, with their kinase domain exhibiting 92% homology (Nakagawa et al., 1996). Despite their molecular similarity, numerous studies have revealed differences in the distribution and actions of these two kinases (Hartmann et al., 2015). Although the expression of ROCK2 in the hypothalamus in normal mice is more prominent than that of ROCK1 (not shown), we found that only ROCK1 is implicated in the regulation of energy balance via MC4R-expressing neurons. Combined with our previous findings that ROCK1 in hypothalamic POMC and AgRP neurons is necessary for the homeostatic control of leptin-mediated energy metabolism (Huang et al., 2012), it is likely that ROCK1 is an indispensable player for the metabolic actions of leptin and melanocortin.

In conclusion, our work identifies ROCK1 as a novel regulator of melanocortin action on food consumption and establishes a new MC4R→Gα_12_→ROCK1→AMPK signaling axis for melanocortin’s anorexigenic action (Fig. 8). This model provides a new mechanism for understanding melanocortin’s metabolic action and the pathogenesis of obesity. Specifically targeting ROCK1 in the hypothalamus could provide a new strategy to combat obesity and its related complications.

**Figure 8.**
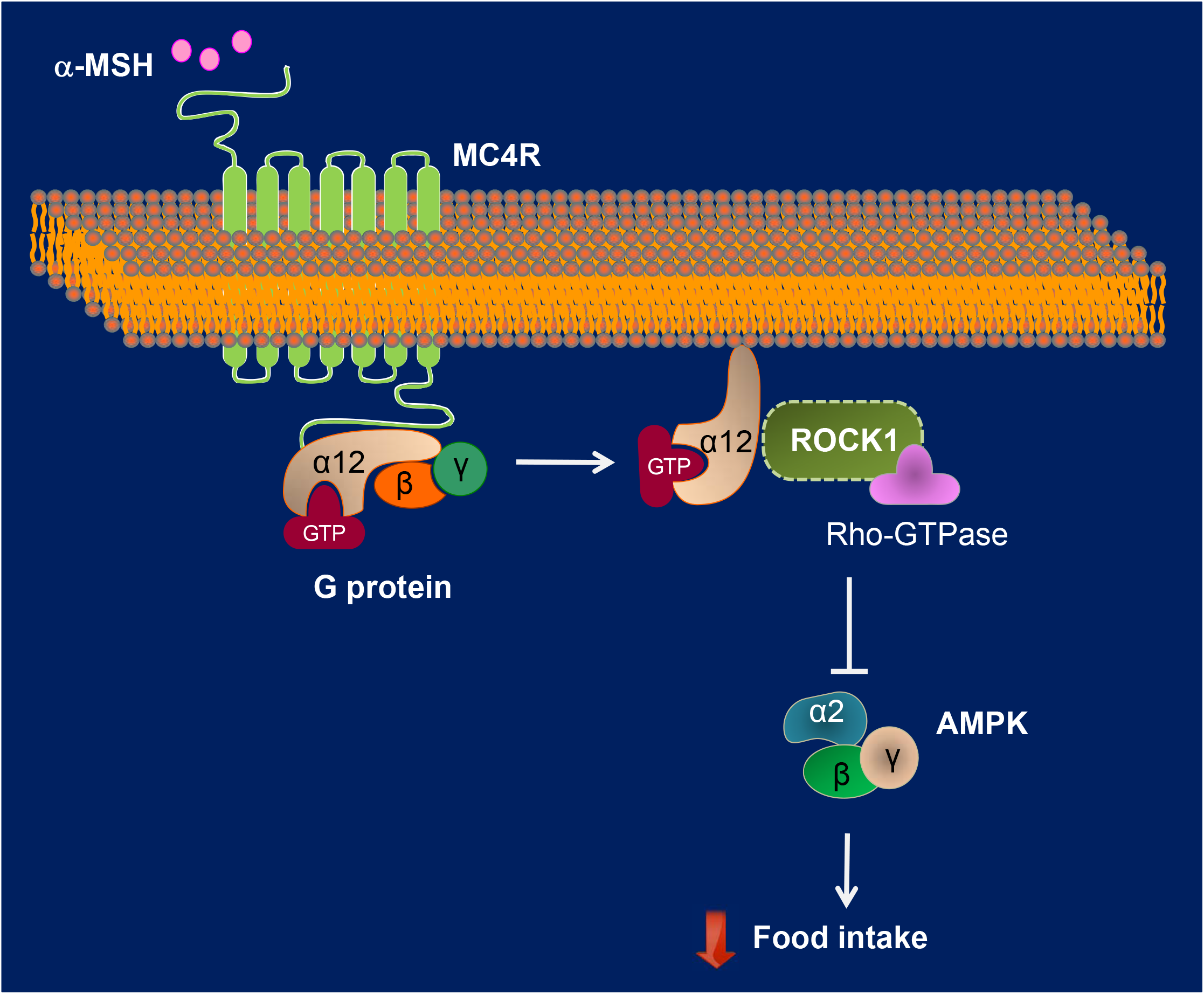
Proposed mechanism for the role of ROCK1 in the anorexigenic action of melanocortin. Upon melanocortin stimulation, the physical interaction of MC4R-Gα_12_ increases in response to α-MSH, and Gα_12_ disassociates from MC4R and then binds to ROCK1. The Gα_12_/ROCK1 complex signals a downstream mediator of AMPK, which is an essential step in melanocortin action in MC4R-expressing neurons. As a result, anorexigenic action is promoted.

## MATERIALs AND METHODs

### Animal care

All animal care and experimental procedures were conducted in accordance with the National Institute of Health’s Guide for the Care and the Use of Laboratory Animals and approved by the Institutional Animal Care and Use Committee of Beth Israel Deaconess Medical Center. Mice were allowed to access standard chow (Teklad F6 Rodent Diet 8664, Harlan Teklad) and water provided *ad libitum* and were housed at 22–24 °C with a 12 h light-dark cycle with the light cycle starting from 6:00 a.m. to 6:00 p.m. Animals were individually housed for studies of food intake, locomotor activity, energy expenditure and after stereotactic surgery. Otherwise, mice were group housed.

### Generation of MC4R-expressing neuron-specific ROCK1 and ROCK2-deficient mice

Mice bearing a *loxP*-flanked ROCK1 allele (ROCK1^*loxP/loxP*^) were generated and maintained as previously described (Garfield et al., 2015, Huang et al., 2012). Mice lacking ROCK1 in MC4R-expressing neurons (MC4R-Cre; ROCK1^*loxP/loxP*^) were generated by mating ROCK1^*loxP/loxP*^ mice with MC4R-2A-Cre transgenic mice (gift from Dr. Brad Lowell, Beth Israel Deaconess Medical Center, Boston, MA). Mice bearing a *loxP*-flanked ROCK2 (ROCK2^*loxP/+*^) were generated by Ingenious Targeting Laboratory (Stony Brook, NY). Briefly, a BAC clone (C57BL/6, RP:23 173D24 clone) containing a 9.4 kb fragment of ROCK2 genomic DNA was used to generate a targeting vector. Five independent ROCK2^flox/+^ ES clones were identified, which were injected into C57BL/6 blastocysts to generate chimeric mice. The chimeric mice were bred with wild type C57BL/6 mice for germline transmission. Heterozygous animals were then crossed with mice expressing flpe-recombinase in the germline (Flipper mice, from The Jackson Laboratory) to delete the FRT-flanked Neo cassette. Offspring of these mice were heterozygous for the desired ROCK2^flox/+^ allele. MC4R-Cre: ROCK2^*loxP/loxP*^ mice were generated by mating ROCK2^*loxP/loxP*^ mice with MC4R-2A-Cre transgenic mice. All experimental mice were on a mixed background.

### Body composition and food intake measurements

Mice were weighted from 4 weeks of birth and weekly thereafter. Fat and lean body mass were assessed by EchoMRI (Echo Medical Systems, Houston, TX). Fat pads were harvested and weighed. For the measurement of daily food intake, mice were individually housed for 1 week prior to the measurement of food intake. Food intake was then measured over a 7-day period. Food intake data for 7 days were combined, averaged, and analyzed by the unpaired Student t-test. For the analysis of food intake after fasting, after a 1 week acclimatization period, mice were then acclimatized to feeding from 3 pellets (maximum ~9g) every day for 3 days. Mice were then fasted overnight and the food intake from 3 pellets (maximum ~9g) was measured at 1, 8 and 24 h. Food intake data were analyzed as above. To assess the exact amount of food intake, a white bedding paper under the food bowl was used to collect food waste during the course of the food intake measurements. The uneaten food was collected and measured. This amount was excluded from the total amount of food intake. For α-MSH-induced and MTII-induced food intake, all animals were single housed for at least 1 week following surgery and handled for 7 days before the assays to reduce the stress response. Experimental mice were fasted for overnight. Food intake was measured over 24 hours after the injection of α-MSH (2 nmol/2 μl, ICV), MTII (5mg/kg, IP) or the vehicle.

### Blood parameter measurements

Blood was collected either from random-fed or overnight fasted mice via the tail. Blood glucose was measured using a OneTouch Ultra glucose meter (LifeScan, Inc., Milpitas, CA). Serum insulin and leptin levels were measured by ELISA (Crystal Chem, Chicago, IL). Serum total cholesterol and triglyceride levels were determined by enzymatic methods (StanbioLaboratory, Boerne, TX). The range and sensitivity for these parameters are: glucose, 40–500 mg/dL; leptin, 1–25.6 ng/mL (sensitivity, 200 pg/mL using a 5-μL sample); insulin, 0.1–12.8 ng/mL (sensitivity, 0.1 ng/mL using a 5-μL sample); total cholesterol, 1–750 mg/dL; and triglyceride, 1–1000 mg/dL.

### Energy expenditure, respiratory exchange ratio and locomotor activity

Energy expenditure was measured by assessing oxygen consumption with indirect calorimetry. Individually housed male mice maintained on a chow diet until 16 weeks of age were studied using the Comprehensive Lab Animal Monitoring System (CLAMS, Columbus Instruments, Columbus, OH). Mice were acclimated in the CLAMS chambers for 72 hours before data collection. Mice had free access to food and water for the duration of the studies. During the course of the energy metabolism measurements (O_2_ and CO_2_) using CLAMS, high variations (overlapping) in measurements emerged at individual time points between groups. This did not allow us to statistically analyze individual time points. To enhance the statistical power of these measurements, we combined each value from the individual time points and analyzed the data by unpaired Student’s *t* tests to compare the two groups.

### Glucose tolerance and insulin tolerance test

For the glucose-tolerance test (GTT), mice were fasted overnight, and blood glucose was measured 0, 15, 30, 60, 90 and 120 minutes after an intraperitoneal injection of glucose (1.0 g/kg of body weight). For the insulin tolerance test (ITT), food was removed for 4 hours in the morning, and blood glucose was measured 0, 15, 30, 60, 90 and 120 minutes after an intraperitoneal injection of human insulin (0.75 IU/kg of body weight; Humulin R, Eli Lilly). The area under the curve or above the curve for glucose was calculated using the trapezoidal rule for GTT or ITT (Lee et al., 2009).

### Quantitative Real-Time PCR

Total RNA was extracted from each tissue or cell using a TRIzol reagent (Invitrogen, CA) and subjected to quantitative real-time PCR as described (Lee et al., 2014). Single stranded cDNA from the total RNA was synthesized with a RT-PCR kit (Clontech, Mountain View, CA) according to the kit’s instructions. qRT-PCR was performed with an Applied Biosystems 7900HT Fast System using a SYBR Green PCR Mastermix reagent (Applied Biosystems, Foster City, CA). Relative mRNA expression levels were determined using the 2^-ΔΔCT^ method normalized to 36B4. Gene-specific primer sequences are listed in supplementary Table 1.

### Cell culture

GT1-7 mouse hypothalamic GnRH neuronal cells (Merck Millipore, Burlington, MA) were maintained in high glucose DMEM (Thermo fisher scientific, Waltham, MA) supplemented with 10% FBS (Thermo fisher scientific) and 1% Antibiotic-Antimycotic (Thermo fisher scientific). Platinum-E retroviral packaging cells (PLAT-E, Cell Biolabs, Inc., San Diego, CA) were maintained in high glucose DMEM supplemented with 10% FBS, 1% Antibiotic-Antimycotic, Blasticidin (10 μg/ml, Thermo fisher scientific), and Puromycin (5 μg/ml, Thermo fisher scientific).

### Generation of GT1-7 cell line overexpressing MC4R-3×HA (GT1-7_MC4R_)

For the generation of the retroviral vector encoding MC4R-3×HA, the cDNA encoding human MC4R-3×HA was amplified from pcDNA3.1-MC4R-3×HA (cDNA resource center, Bloomsberg, PA). The PCR product was cloned into BamHI and XhoI sites of pMXs-IRES-Bsd retroviral expression vector (Cell Biolabs, Inc.). The primer sequences were 5’-AAAA AGGATCCACTTAAGCTTGGTACCAC-3’ and 5’-TCTAGACTCGAGTTAATATCTGCT AGACA-3’. Retroviral particles were generated by transfection of PLAT-E cells (Cell Biolabs, Inc.) with the retroviral vector encoding MC4R-3×HA using Lipofectamine 3000 (Thermo fisher scientific), in accordance with the manufacturer’s instruction. GT1-7 cells were infected with the retrovirus encoding MC4R-3×HA using ViraDuctin^™^ Transduction Kit (Cell Biolabs, Inc.) according to the manufacturer’s protocol. After infecting for 24 h, the cells were cultured in the presence of Blasticidin (10 μg/ml, Thermo fisher scientific). Blasticidin-resistant cells were selected for 2 weeks and used for further study. The overexpression of MC4R-3×HA was verified by immunoblotting or immuno-fluorescent staining.

### Intracellular cAMP measurements

GT1-7_MC4R_ cells were incubated for 24 h on 96-well plates. The cells were serum-starved for 4 h, and then changed to an induction buffer (serum-free media containing 500 μM isobutyl-1-methylxanthine (IBMX, Sigma-Aldrich, St. Louis, MO) and 100 μM 4-(3-butoxy-4-methoxybenzyl) imidazolidone (Ro 20-1724, Sigma-Aldrich)). The cells were pre-treated with SHU9119 (1, 5, and 10 μM) or fasudil (1, 10, and 30 μM) for 1 h, and then stimulated with α-MSH (1 μM) for 15 min. Intracellular cAMP levels were measured using cAMP-Glo assay kit (Promega, Madison, WI) according to the manufacturer’s instruction and quantified on a plate-reading luminometer.

### Proximal ligation assay (PLA)

GT1-7_MC4R_ cells were plated on a 8-well chamber slide (Thermo fisher scientific) overnight. The cells were serum-starved for 4 h and pre-treated with fasudil (30 μM) or SHU9119 (10 μM) for 1 h and then treated with α-MSH (1 μM) for 15 min. The cells were washed 3 times with ice-cold PBS and fixed with methanol. Cells were incubated with primary antibodies to anti-rabbit ROCK1 (sc-5560, 1:100, Santa Cruz Biotechnology), anti-mouse HA (H3663, 1:100, Sigma-Aldrich), anti-rabbit HA (H6908, 1:100, Sigma-Aldrich), anti-mouse Gα_12_ (sc-515445, 1:100, Santa Cruz Biotechnology), or anti-mouse Gαs (sc-135914, 1:100, Santa Cruz Biotechnology) overnight. PLA was performed using PLA PLUS or MINUS probes for rabbit or mouse anti-serum and Duolink In Situ Detection Reagents Red (Sigma-Aldrich, St. Louis, MO) according to the manufacturer’s instruction. The chamber slide was mounted using Duolink In Situ Mounting Medium with DAPI (Sigma-Aldrich) and was viewed under a laser confocal fluorescence microscope (LSM-710, Carl Zeiss). Red fluorescent signal intensity was automatically determined by using Image J software (National Institutes of Health, Rockville, MD) and quantified by calculating the red fluorescent signal intensity with the indicated number of nuclei.

### Immunofluorescent analysis

GT1-7_MC4R_ cells were fixed in methanol and rinsed with phosphate-buffered saline. The fixed cells were incubated with an antibody against HA antibody (H6908, 1:100) at 4°C overnight. Cells were washed and incubated for 1h with Alexa Fluor 488-conjuated antibody to rabbit IgG (A-21206, 1:100, Thermo fisher scientific). Images of green fluorescence were assessed by laser confocal fluorescence microscopy.

### Immunoblotting analysis

Membrane proteins and cytosolic proteins were isolated using Mem-PER^™^ Plus membrane protein extraction kit (Thermo fisher scientific) according to the manufacturer’s instruction. Tissues were homogenized in lysis buffer as described previously (Furukawa et al., 2005). Cell and tissue lysate proteins (20–50 μg of protein) were resolved by SDS-PAGE and transferred to nitrocellulose membranes. The membranes were incubated with an antibody against HA (H6908, 1:10,000). The bands were visualized with enhanced chemiluminescence and quantified by densitometry.

### ROCK1 activity and α2 AMPK activity assay

Male ROCK1^*loxP/loxP*^ (control) and MC4R Cre; ROCK1^*loxP/loxP*^ mice (12 weeks of age) fed a normal chow diet were fasted overnight. Mice were administered intracerebroventricular (ICV) injections of α–MSH (2 nmol/2μl) or intraperitoneal (IP) injections of MTII (5mg/kg) or vehicle, and scarified 30 minutes following administration for ICV injections or 45 minutes for IP injections. The hypothalamus was rapidly removed for measurement of enzymatic ROCK1 and α2 AMPK activity. GT1-7_MC4R_ cells were starved for 3 hours and stimulated with α-MSH (1 μM) in the presence of fasudil (30 μM) for 15 min, and harvested for measurements of enzymatic ROCK1 and α2 AMPK activity. For ROCK1 activity, lysates from the mouse hypothalamus (75 μg of protein) or GT1-7_MC4R_ cells (50 μg of protein) were subjected to immunoprecipitation overnight with 10 μl of a polyclonal ROCK1 antibody (sc-6055, Santa Cruz Biotechnology) coupled to protein G-Sepharose beads (GE healthcare, Piscataway, NJ). Immune pellets were washed and resuspended in 50 μl of kinase mixture [20mM Tris (pH 7.5), 5 mM MgCl_2_, 100 mM KCl, 0.1 mM DTT, 100 μM ATP, 1 mM EDTA, 1 μM microcystin-LR, 50 μM long S6K substrate peptide (Millipore) and 1 μCi (γ-^32^P) ATP] and incubated at 30°C for 30 min. Samples (40 μl) were spotted onto phosphocellulose p81 paper (Whatman) and washed with 75 mM orthophosphoric acid. Bound radioactivity was measured by scintillation (Huang et al., 2013b). For α2 AMPK activity, lysates from mice hypothalami (75 μg protein) or GT1-7_MC4R_ cells (50 μg of protein) were subjected to immunoprecipitation overnight with 1 μg of a specific α2 AMPK antibody (ab3760, Abcam, Cambridge, MA) coupled with protein G-Sepharose beads (GE Healthcare, Piscataway, NJ). Immune pellets were washed and resuspended in 50 μl of kinase mixture [833 μM DTT, 5 mM MgCl2, 200 μM ATP, 200 μM AMP, 100 μM SAMS (substrate for AMP-activated Protein Kinase, Millipore), 0.2 μCi (γ-^32^P) ATP] and incubated at 30°C for 30 min. Samples (40 μl) were spotted onto phosphocellulose p81 paper (Whatman) and washed with 75 mM orthophosphoric acid. Bound radioactivity was measured by scintillation (Huang et al., 2018).

### Statistical analysis

Data are presented as mean ± SEM. Differences between groups were assessed for statistical significance by either the unpaired Student’s *t* test or ANOVA when appropriate. Differences were considered significant at *p* < 0.05. Statistical analyses were performed using Prism 5.0 software (GraphPad Software, La Jolla, CA).

## AUTHOR CONTRIBUTIONS

Y.B.K., S.S.K., W.M.H. and W.M.Y. designed the study. S.S.K., and W.M.H. performed most of the experiments with MC4R-neuron-specific ROCK1-deficient and ROCK2-deficient mice. S.S.K., W.M.H. and H.L. bred and maintained the experimental mice and genotyped the mice. W.M.Y. and K.S.P. generated the GT1-7 cell line overexpressing MC4R-3XHA and performed *in vitro* experiments. S.S.K., W.M.H and Y.D. measured ROCK1 and AMPK activity. All authors analyzed and interpreted the experimental data. SSK, H.L., and YBK wrote the manuscript.

## ACKNOWLEDGMENTS

This work was supported by grants from the National Institutes of Health (R01DK083567 to Y.B.K.) and the National Research Foundation of Korea the National Research Foundation of Korea (2018R1C1B6002854 to S.S.K). We thank Brad Lowell for MC4R-2A-Cre transgenic mice; Sung Man Cho, Min Cheol Kang and John Campbell for technical help; and Ya-Xiong Tao, Qingchun Tong, and Brad Lowell for their helpful discussions.

