## Supplemental data for "Rho-kinase mediates the anorexigenic action of melanocortin by suppressing AMPK"

**Supplemental Figure Legends**

**Supplemental Figure 1. Deletion of ROCK1 in MC4R-expressing neurons causes insulin resistance and glucose intolerance.**

(A) ITT was performed 4 h after food removal in 16-week-old male ROCK1$^{loxP/loxP}$ (control) and MC4R-Cre; ROCK1$^{loxP/loxP}$ mice (n = 6–7). Mice were injected with insulin (IP 0.75 U/kg of body weight). Blood glucose was determined by glucometer at the indicated times. The area above the curve for glucose was calculated using the trapezoidal rule for ITT. Data are presented as means ± SEM. *p < 0.05 vs. control mice

(B) GTT was performed on 16-week-old male ROCK1$^{loxP/loxP}$ (control) and MC4R-Cre; ROCK1$^{loxP/loxP}$ mice (n = 6–7) after overnight fasting. Mice were injected with glucose (IP 1.0g/kg of body weight). The area under the curve for glucose was calculated using the trapezoidal rule for GTT. Blood glucose was determined at the indicated times by glucometer. Data are presented as means ± SEM. *p < 0.05 vs. control mice

**Supplemental Figure 2. Respiratory exchange ratio and locomotor activity in MC4R-neuron-specific ROCK1 deficient mice.**

(A) The average hourly respiratory exchange ratio (RER) corresponding light and dark phase RER (12 h average) in male 16-week-old ROCK1$^{loxP/loxP}$ (control) and MC4R-Cre; ROCK1$^{loxP/loxP}$ mice (n = 8) are shown. RER is calculated from the ratio between $CO_2$ and $O_2$. Data are presented as means ± SEM.

(B) The average hourly locomotor activity and corresponding light and dark phase locomotor activity (12 h average) in male ROCK1$^{loxP/loxP}$ (control) and MC4R-Cre; ROCK1$^{loxP/loxP}$ mice (n = 8) are shown. Locomotor activity was measured by CLAMS. Data are presented as means ± SEM. $^{*}P < 0.05$, vs. control

**Supplemental Figure 3. Hypothalamic mRNA expression of neuropeptides in MC4R-neuron-specific ROCK1 deficient mice.**

mRNA expressions of POMC, AgRP, and NPY were assessed by real time RT-PCR in male 16-week-old ROCK1$^{loxP/loxP}$ (control) and MC4R-Cre; ROCK1$^{loxP/loxP}$ mice (n = 6–7). Data are presented as means ± SEM.

**Supplemental Figure 4. Generation of ROCK2*loxP/loxP* mice.**

(A) Schematic representation of the wild type ROCK2 allele, targeting vector, targeted allele and flox allele. The targeting vector containing *loxP* sites flanking exon 3 of ROCK2 was injected into embryonic stem cells.

(B) Southern blot analysis of G418 resistant ES cells. DNA encoding the ROCK2 targeting vector was linearized by Not I and then transfected by electroporation of iTL BA1(C57BL/6 x 129/SvEv) hybrid embryonic stem cells. After selection with G418 antibiotic, surviving clones were expanded for Southern analysis to identify recombinant ES clones. DNA was digested with BamH I. Blotting of ES cell DNA shows the WT C57BL/6/129 allele, a band of 15.2 kb, and the ROCK2 targeted C57BL/6 allele which appears as a band shifted from 15.2 to 13.1 kb.

(C) Genotyping of ROCK2*loxP/loxP* mice. Tail DNA samples from F1 mice were amplified by PCR using genomic primer pair, which flanks one of loxP sites. With the addition of the *loxP* site, the amplified product is 363 bp. The amplified product is 318 bp for wild-type (or without a loxP site). The positive control (positive ES clone) is indicated by (+).

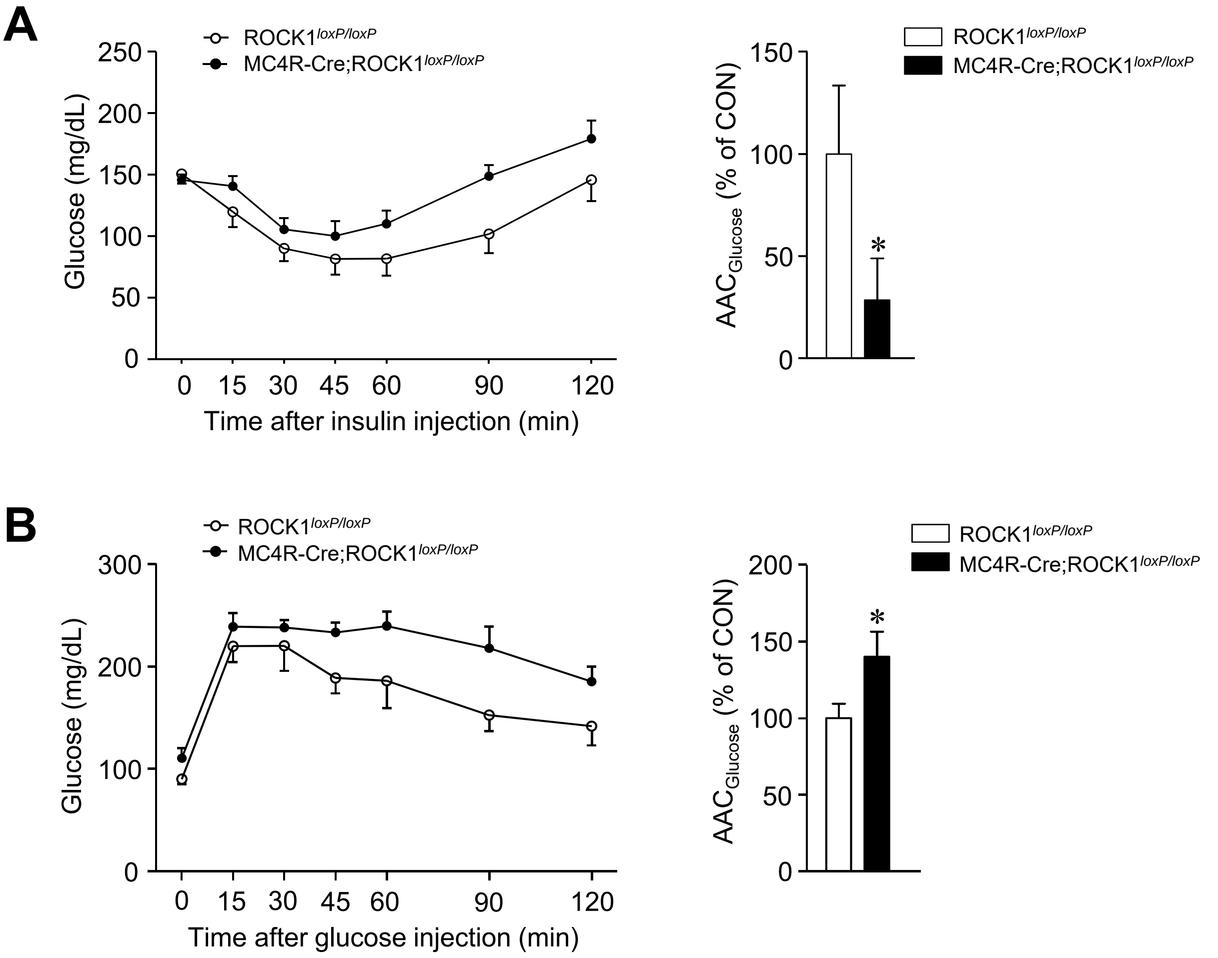


Supplementary Fig. 1

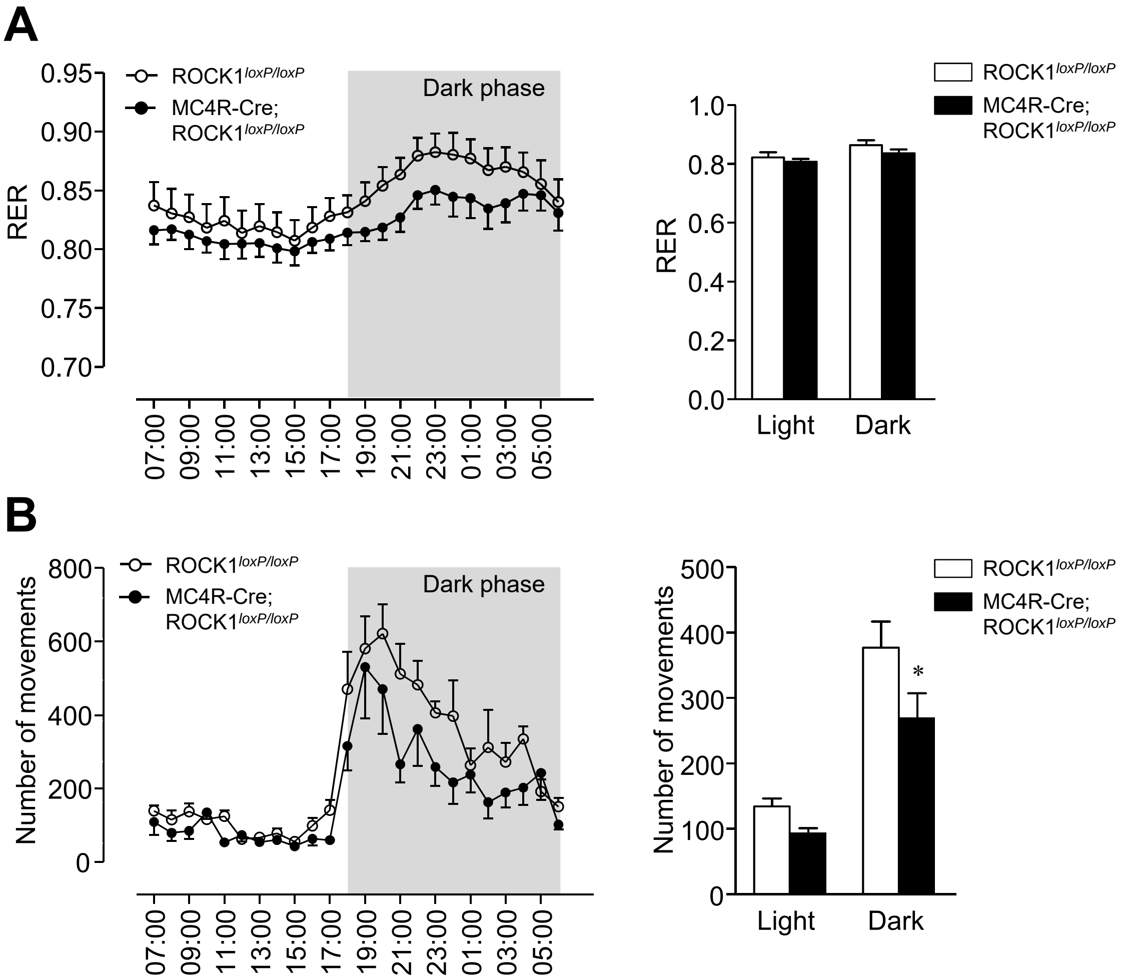


Supplemental Fig. 2

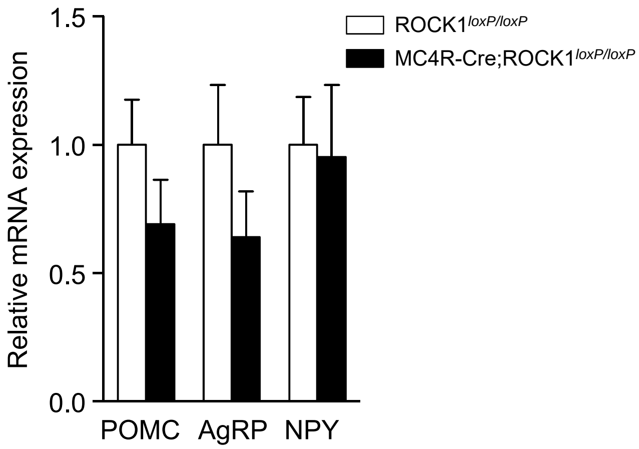


Supplemental Fig. 3

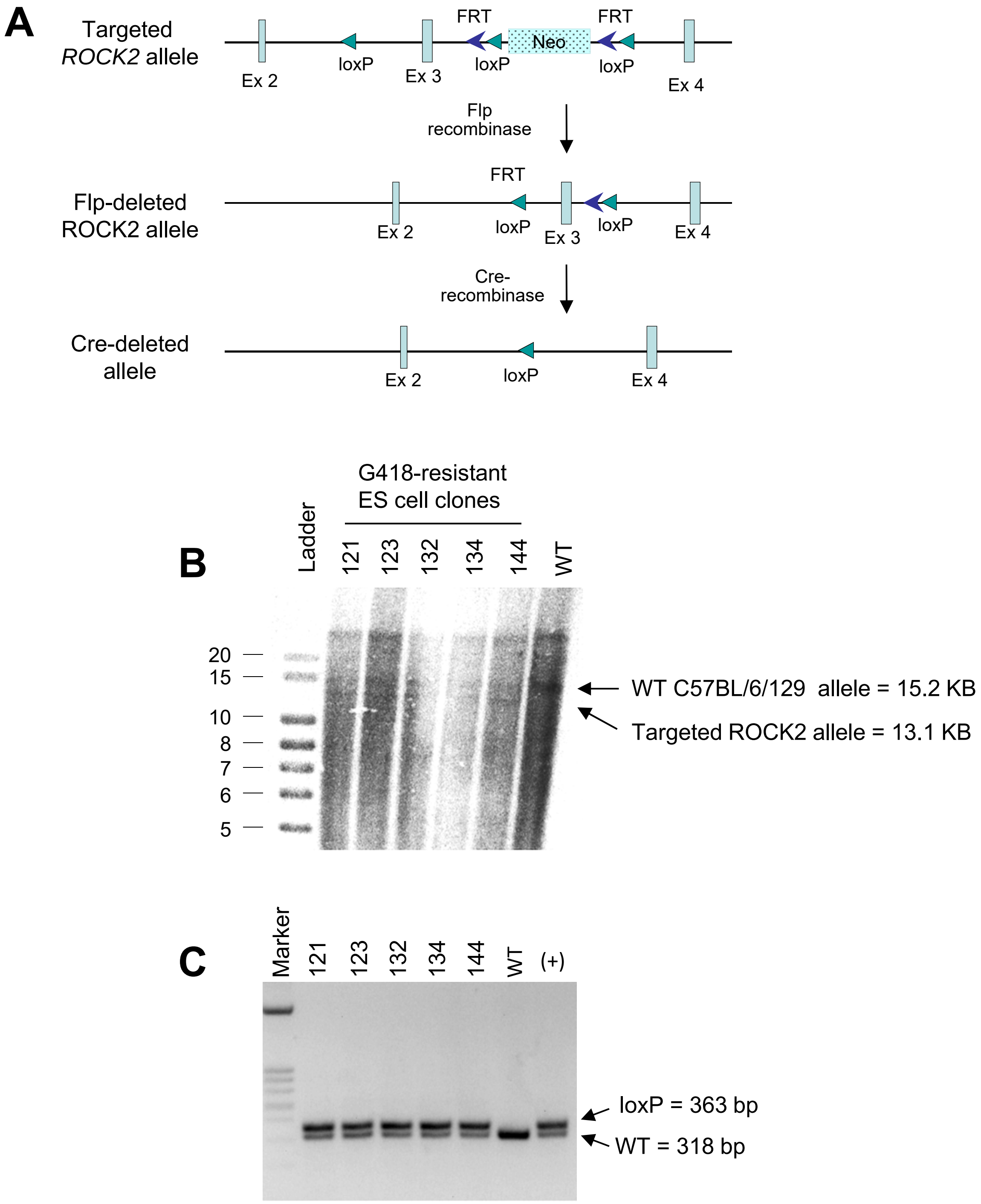


Supplemental Fig. 4

**Supplementary Table 1. Primer sequences used in qRT-PCR**

| **Gene** | **Primer** | **Sequence** |
|---|---|---|
| 36B4 | Forward | AGATGCAGCAGATCCGCAT |
| | Reverse | GTTCTTGCCCATCAGCACC |
| Pgc1α | Forward | CCCTGCCATTGTTAAGACC |
| | Reverse | TGCTGCTGTTCCTGTTTTC |
| Ucp1 | Forward | ACTGCCACACCTCCAGTCATT |
| | Reverse | CTTTGCCTCACTCAGGATTGG |
| *Cox8b* | Forward | GAACCATGAAGCCAACGACT |
| | Reverse | GCGAAGTTCACAGTGGTTCC |
| Cox7a1 | Forward | GCTCTGGTCCGGTCTTTTAG |
| | Reverse | CTTTCAAGTGTACTGGGAGGTC |
| Dio2 | Forward | GTGGCTGACTTCCTGTTGGT |
| | Reverse | GCACACACGTTCAAAGGCTA |
| Elovl3 | Forward | TCCGCGTTCTCATGTAGGTCT |
| | Reverse | GGACCTGATGCAACCCTATGA |
| Cpt1b | Forward | TCTTGCAGTCGACTCACCTT |
| | Reverse | TCCACAGGACACATAGTCAGG |
